# A Mathematical Model which Examines Age-Related Stochastic Fluctuations in DNA Maintenance Methylation

**DOI:** 10.1101/2021.02.05.429896

**Authors:** Loukas Zagkos, Jason Roberts, Mark Mc Auley

## Abstract

Due to its complexity and its ubiquitous nature the ageing process remains an enduring biological puzzle. Many molecular mechanisms and biochemical process have become synonymous with ageing. However, recent findings have pinpointed epigenetics as having a key role in ageing and healthspan. In particular age related changes to DNA methylation offer the possibility of monitoring the trajectory of biological ageing and could even be used to predict the onset of diseases such as cancer, Alzheimer’s disease and cardiovascular disease. At the molecular level emerging evidence strongly suggests the regulatory processes which govern DNA methylation are subject to intracellular stochasticity. It is challenging to fully understand the impact of stochasticity on DNA methylation levels at the molecular level experimentally. An ideal solution is to use mathematical models to capture the essence of the stochasticity and its outcomes. In this paper we present a novel stochastic model which accounts for specific methylation levels within a gene promoter. We quantify the uncertainty of the eventual cite-specific methylation levels for different values of methylation age, depending on the initial methylation levels. Our model predicts the observed bistable levels in CpG islands. In addition, simulations with various levels of noise indicate that uncertainty predominantly spreads through the hypermethylated region of stability, especially for large values of input noise. A key outcome of the model is that CpG islands with intermediate methylation levels tend to be more susceptible to dramatic DNA methylation changes towards both hypomethylation and hypermethylation, due to increasing methylation age.

## 1 Introduction

DNA methylation is central to the regulation of normal gene expression. However, during ageing it has been found in a wide variety of tissues that the methylation status of the genome alters [El Khoury et al., 2019] [Yu et al., 2019]. This phenomenon is characterised by global genome hypomethylation and site specific hypermethylation [McLain and Faulk, 2018]. The age related epigenomic transformation of the genome has key implications for healthspan because it is associated with diseases such as cancer, Alzheimer’s disease, cardiovascular disease, osteoporosis and diabetes [Semick et al., 2019] [Davegardh et al., 2018]. Moreover, there is strong and intriguing experimental evidence which suggests that DNA methylation changes could be used to track the trajectory of the ageing process [Xie et al., 2019]. This derives from the remarkable studies which have accurately managed to estimate chronological age based on several hundred sites specific alterations to the methylome [Bell et al., 2019][Horvath and Raj, 2018]. These changes are thought to be the result of the diminishing capacity of an intrinsic epigenetic maintenance system [Horvath, 2013][Horvath et al., 2016]. Collectively, such findings provide compelling evidence that age related changes to the molecular reactions which are responsible for regulating DNA methylation could impact our risk of disease and possibly even be arbiters of the ageing process itself [Ciccarone et al., 2018] [Morgan et al, 2018].

DNA methylation reactions take place mainly at CpG dyads, where the methyl group is covalently bonded to the fifth carbon of the cytosine at the CpG site. A CpG site is a 5’-3’ Cytosine – Guanine dinucleotide sequence within the DNA molecule with “p” being the phosphate group between the two nucleotides. A dyad incorporates two CpG sites, one on each strand of DNA, while regions of DNA which have a high occurrence of CpG sites are known as CpG islands (CGIs). These areas are made up of 500-2000 primary unmethylated base pairs which cover approximately one percent of the human genome [Antequera and Bird, 2018]. Although there is not a wealth of CGIs, they have a fundamental role to play because their methylation status is synonymous with gene promoter activity [Morgan and Marioni, 2018]. In particular, hypermethylation, of CGIs commonly accompanies the transcriptional silencing of gene promoters; a feature which is commonplace in diseases such as cancer [Skvortsova et al., 2019]. In addition to pathologies such as cancer, increasing age has been correlated with the hypermethylation of a range of gene promoters [Guarasci et al., 2018]. Thus, it is possible ageing effects the molecular machinery which underpins the regulation of DNA methylation.

In recent years mathematical models have played a vital role in improving our understanding of the dynamics of DNA methylation. A number of these models are reviewed in [Mc Auley et al., 2018]. The mechanisms which govern DNA methylation have been modelled using two different mathematical approaches. Firstly, they have been modelled using a deterministic framework. Otto and Walbot were the first to present a mathematical model to describe the characteristic features of DNA methylation [Otto and Walbot, 1990]. The authors assumed a recursive model of methylation that represents the kinetics between de novo and maintenance methylation and demethylation. Following their model, Pfeifer and colleagues attempted to depict the methylation patterns in a growing population, by developing a set of two deterministic differential equations to specify the level of methylation in a population of growing cells [Pfeifer et al., 1990]. This deterministic approach was adopted also by McGovern and colleagues to represent the reactions associated with DNA methylation when they used partial differential equations to model this system [McGovern et al., 2012]. Finally, in recently published work [Zagkos et al., 2019] we described the construction and examination of linear and nonlinear deterministic models of DNA methylation in order to gain insights into the connection between the dynamics of DNA methylation and health.

Although all the models mentioned thus far have improved our understanding of DNA methylation, they fail to take account of the fact that living systems are inherently noisy and are optimized to function in the presence of stochastic fluctuations [McAdams and Arkin, 1999]. For example, cells are noisy biochemical reactors [Thattai and van Oudenaarden, 2001]. Likewise, the process of DNA methylation is considered to be noisy [Singer et al., 2014]. By the term “noise” we mean the variability that typically arises from the inherent stochastic nature of the individual biochemical processes associated with DNA methylation.

Clearly it is cogent to represent stochasticity within models which depict cellular process such as DNA methylation and there are valid experimental reasons for doing this. It is known the binding and diffusion dynamics of molecules generates noise, and it is highly probable DNA methylation is subject to this. In fact, experimental evidence supports this idea as methylation fidelity studies suggest the DNA methylation dynamics are characterised by stochasticity [Landan et al., 2012]. Moreover, the notion that there is a probabilistic underpinning to DNA methylation has been suggested by [Jeltsch and Jurkowska, 2014]. Consequently, a stochastic approach to modelling DNA methylation has been adopted by [Haerter et al., 2014] who utilised the Gillespie algorithm [Gillespie 1976] to model the dynamics of this process. Others have made efforts to represent the hypothesis that DNA methylation levels are heavily dependent on density of a CpG cluster. The conjecture was made that spatial dependent collaboration between CpG sites is a key factor. This notion was consolidated by the idea that methylated CpG sites care capable of sequestering enzymes that interact with CpG sites in the neighbouring area [Dodd et al., 2007]. Additionally, it was assumed that unmethylated CpGs can recruit demethylation enzymes which predispose neighbouring CpGs to demethylation [Loevkvist et al., 2016], where the authors used a standard Gillespie algorithm to update the state of the CpG sites. Recently, a stochastic model was proposed for the dynamics of the post-replicative restoration of methylation patterns [Utsey and Keener, 2020]. The model considers the recruitment of Dnmts and demethylating enzymes to regions of hypermethylation and hypomethylation, respectively, and also includes the interaction between Dnmt1 and PCNA, an enzyme that localizes Dnmt1 to the replication complex.

There are many different types of stochastic methods which can be used to examine biological phenomena at the molecular level [Tsimring, 2014]. For instance, the Gillespie algorithm has been applied ubiquitously [Mc Auley et al., 2017]. Although it is a powerful tool which is statistically exact, it is computationally intensive [Szekely and Burrage, 2014]. Moreover, modelling with stochastic differential equations has been successfully applied to an array of cellular processes [Saarinen et al., 2006] [Lei and Mackey, 2007] [Saarinen et al., 2008] [Dangerfield et al., 2012]. Consequently, in this work we introduce randomness to the parameters of our recently published model [Zagkos et al., 2019] in the form of ‘white noise’ in order to account for the uncertainty in the parameter values due to the noisy molecular processes which underpin DNA methylation. Here, ‘white noise’ is considered to be the generalised derivative of the standard Wiener process, as shown in Appendix A. A standard Wiener process (see Appendix A) is a convenient way to mathematically define and quantify randomness in mathematical models. This enables a new stochastic model of DNA methylation to be developed. We use this model to investigate how age-associated maintenance methylation noise influences gene promoter methylation; and reveal a novel way to identify regions of DNA methylation instability within gene promoters. We conclude by discussing the implications of our findings for understanding health and ageing.

## 2 Models and Methods

Recently we introduced a new mathematical model of DNA methylation [Zagkos et al., 2019]. This deterministic model captures the bistable methylation states which have been observed experimentally in gene promoters [Bennett and Hasty, 2007].

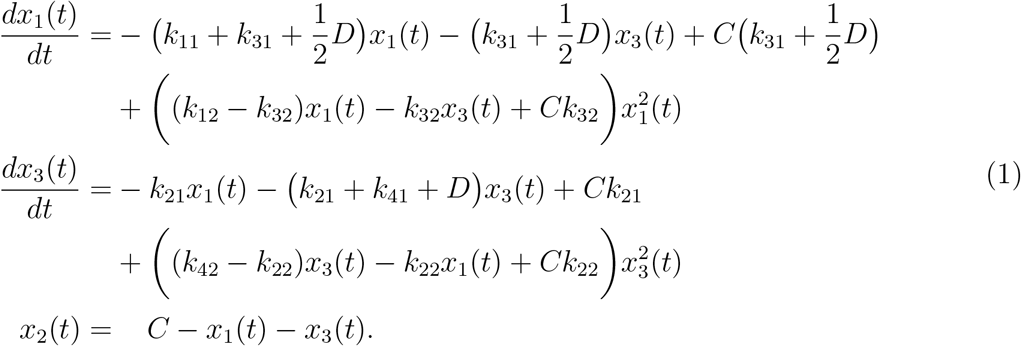

As can be observed in Figure 1, transition rates are considered as functions of the different states of the CpG dyads. *k*_1_ is the methylation rate of the unmethylated CpG dyads *x*_1_, *k*_2_ denotes the methylation rate of the hemimethylated CpG dyads *x*_2_, *k*_3_ is the demethylation rate of the hemimethylated CpG dyads *x*_3_ and *k*_4_ denotes the demethylation rate of the methylated CpG dyads *x*_4_. *D* is the division cell rate and *C* the total number of CpG dyads in the investigation of interest.

**Figure 1:**
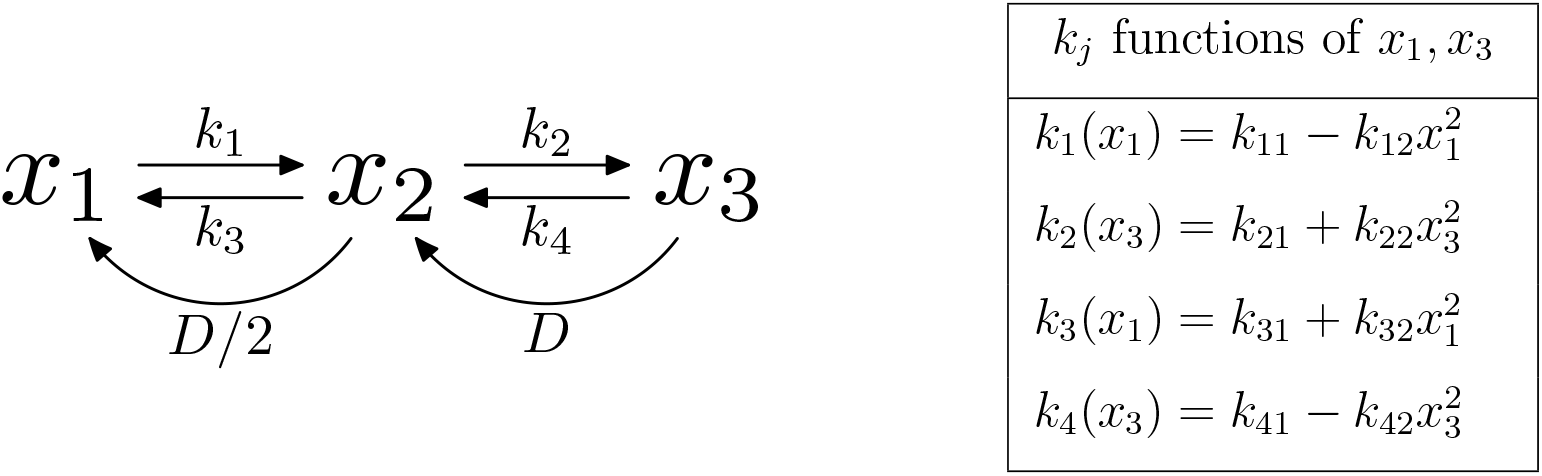
Schematic overview of the model and its transition rates *k*_*j*_, *j* = 1, 2, 3, 4. All parameters *k*_*ij*_ are positive.

Based on the sensitivity analysis originally performed for (1); in descending order the most sensitive parameters of the nonlinear model are: *k*_21_, *k*_32_ and *k*_41_. In this context the term ‘most sensitive’ refers to the parameters which, for a given percentage change in their value, have the most significant effect on the percentage change in the CpG dyads populations.

Despite the simplicity of the local sensitivity analysis, this approach failed to explore the entire parameter space, as it did not account for simultaneous parameter variations. Consequently certain behaviour was overlooked by only considering individual parameter fluctuations. Moreover, this approach was unable to identify interactions between input variables. To overcome this, we conducted additional work to explore in greater depth the parameters of our model.

The method we adopted involved applying a recently developed Bayesian algorithm which uses a Transitional Markov Chain Monte Carlo (TMCMC) sampling method [Larson et al., 2019]. This recently published work describes how we used this approach to identify the most uncertain parameters. In descending order, these were: *k*_21_, *k*_12_, *k*_22_ and *k*_41_. Interestingly, the findings of the TMCMC algorithm agree with the original sensitivity analysis that parameter *k*_21_ is the most sensitive (1).

*k*_21_ represents maintenance methylation. The enzyme is primarily responsible is DNA (cytosine-5)-methyltransferase 1 (Dnmt1). The finding that this is the most sensitive parameter is perhaps unsurprising when one considers the findings of recent mathematical modelling [Busto-Moner et al., 2020]. Busto-Moner and colleagues revealed kinetic heterogeneity in post-replication DNA methylation. Therefore, there is a clear basis for examining stochastic fluctuations in this parameter. Moreover, as the focus of this paper is the intersection between healthspan and DNA methylation, there are additional solid biologically reasons why it is important for this work to examine this parameter. Namely, as our focus is the ageing process it is worth considering how the dynamics of this Dnmt1 alters with age. Age related DNA methylation dysregualtion can at least in part be traced to the alteration of the expression of DNMT1 with age [Ciccarone et al., 2016]. Ageing correlates with increased noise in gene expression, and it is likely this is the case with DNMT1 expression [Bahar et al., 2006]. Despite our focus on DNMT1, this does not mean that uncertainty does not lie within the other parameters and it would remiss to think that this is not the case. However, given the key role of Dnmt1 in post-replication DNA methylation it is important to investigate how fluctuations in this key parameter influences the dynamics of the system and the impact this has for ageing and healthspan.

Using the findings from our previous work, in this paper randomness will be introduced to the model in the form of “white noise” in order to account for parameter uncertainty. Since *k*_21_ was identified as the most uncertain parameter, we construct a new stochastic model introducing “white noise” into parameter *k*_21_ and explore the impact on DNA methylation of different noise levels. If we consider:

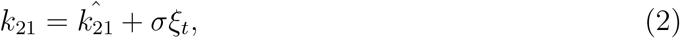

where *σ* > 0 and *ξ_t_* is a white noise process. In this way“randomness” in the form of white noise can be introduced to *k*_21_. Significantly, this parameter represents maintenance methylation. DNMT1 is the enzyme responsible for maintenance methylation. In our previous work parameter 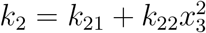. was defined as the transition rate between a hemimethylated and a methylated CpG dyad, namely, the rate of maintenance methylation, which is not constant but varies over time, as a function of CpG dyads. Taking into consideration all the above, equation (2) represents the uncertainty of the enzyme Dnmt1, and thus the variability of maintenance methylation. We substitute expression (2) into system (1) and after several calculations (see Appendix A) we obtain the stochastic system:

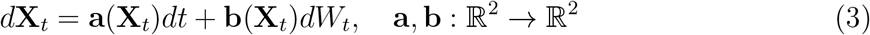

where

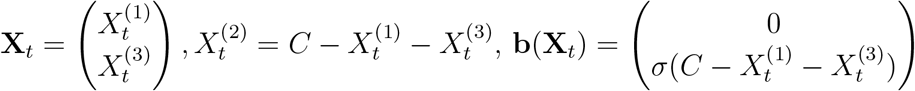

and

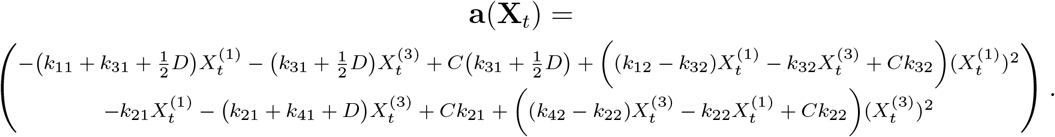

For the value of the parameter 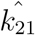, we used the nominal value of the parameter *k*_21_ used in the deterministic model. Thus, for simplicity, we substituted the term 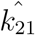 with *k*_21_. 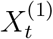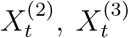 are random variables denoting unmethylated, hemimethylated and methylated CpG dyads, respectively, and *W*_*t*_ is a scalar standard Wiener process.

## 3 Results

We applied the stochastic model (3) to an averaged size CpG cluster of *C* = 100 CpG dyads in total. We considered the following values for the parameters, as outlined previously [Zagkos et al., 2019]: *k*_11_ = 2.1, *k*_12_ = 2 × 10^−5^, *k*_21_ = 10, *k*_22_ = 10^−2^, *k*_31_ = 1, *k*_32_ = 10^−2^, *k*_41_ = 4, *k*_42_ = 2 × 10^−4^. Stochastic trajectories of the solutions of the stochastic system (3) are plotted versus time using Matlab 2017a for the simulations. In this system, noise is added in parameter 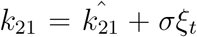. Let us denote *X*^(1)^ = (2.2, 3.1, 94.7)^*T*^, *X*^(2)^ = (93.2, 2, 4.1)^*T*^ and *X*^(3)^ = (33.4, 5.5, 61.1)^*T*^ the three equilibrium points of the nonlinear deterministic system (1). *X*^(1)^ and *X*^(2)^ are stable equilibrium points and *X*^(3)^ is unstable and defines the boundary between the two stable regions. Each of these points are selected as initial values to generate sample paths of the stochastic system (3). For each starting point, we select various values of noise *σ* = 0.01, 0.05, 0.1, 0.5, 1. For each initial value and level of noise we plot two sample paths and the mean of 10^4^ sample paths. The objective is to numerically calculate the values of the equilibrium points and identify their stability. The Euler-Maruyama scheme is used for approximating the solution of the stochastic differential equation (3).

We first set *X*^(1)^ = (2.2, 3.1, 94.7)^*T*^ as the initial value. For *σ* = 0.01, 0.05, 0.1, the sample paths have no fluctuations and look like the deterministic solutions. For *σ* = 0.5 and above, we observe stochastic fluctuations in the sample paths (Supplementary Figure 9). We observe that the noise added to the system influences the stochastic trajectories of *X*_2_ and *X*_3_ significantly more than *X*_1_, when the trajectories remain close to the point *X*^(1)^.

The hemimethylated and methylated CpG dyads fluctuate significantly more than unmethylated. This is because maintenance methylation was considered to be the noisy process. Therefore the hemimethylated and methylated dyads are affected more in this region (where *X*_1_ is small compared to *X*_2_ and *X*_3_). For every noise level up to *σ* = 1, all sample paths converge to the point *X*^(1)^. This can be confirmed by plotting the mean of 10^4^ sample paths over time, which converges to *X*^(1)^, as shown in Supplementary Figure 10.

Next, *X*^(2)^ = (93.2, 2, 4.1)^*T*^ is selected as the initial value. We observe that the sample paths have larger fluctuations near the point *X*^(2)^ than near *X*^(1)^. The sample paths remain in the vicinity of the starting point *X*^(2)^ (Figure 2). If the level of noise is significantly larger than 100%, due to the large oscillations of the sample path, trajectories is more likely to move outside the reasonable space [0*, C*]. This causes the trajectories to become negative or more than the total number *C.* It is possible that this phenomenon can take place for smaller values of noise *σ* due to the stochastic jumps of the Wiener process, but it is more unlikely as the noise decreases [Ditlevsen and Samson, 2013]. Due to the structure of the additive noise (*σ* multiplied by the small number of hemimethylated CpG dyads *X*_2_) it is highly unlikely that any stochastic trajectory could get negative values for input noise values smaller than 100%. Therefore, for meaningful calculations, we are more interested in white noise input values smaller than *σ* = 1. Nevertheless, we plotted values of input noise up to 500% to further investigate the influence of uncertainty in the system (see Supplementary Figure 11).

**Figure 2:**
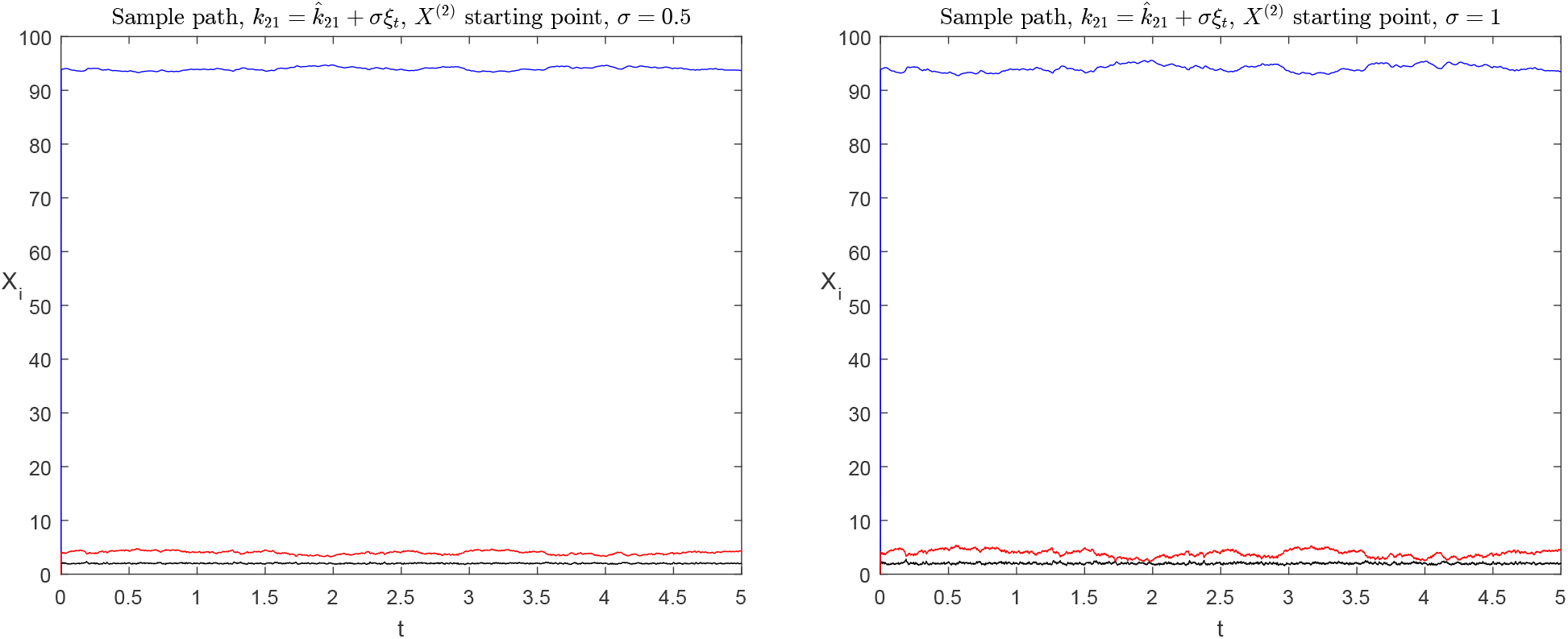
A sample path of (3) with starting point *X*^(2)^ = (93.2, 2, 4.1)^*T*^ with *σ* = 50% and *σ* = 100%. Unmethylated CpG dyads are denoted in blue, hemimethylated in black and methylated in red. The sample paths have more significant fluctuations near the point *X*^(2)^ than near *X*^(1)^. The sample paths remain in the vicinity of the starting point *X*^(2)^.

In contrast to starting points *X*^(1)^ and *X*^(2)^, of particular interest is the case when the starting point is *X*^(3)^ = (33.4, 5.5, 61.1)^*T*^, since it lies at the boundary of the two stability regions. Two kinds of sample paths are obtained when the initial value is *X*^(3)^, as shown in Figures 3, 4, 5 and Supplementary Figure 12. For noise levels greater than *σ* = 0.1, some stochastic trajectories converge to *X*^(1)^ whereas the rest converge to *X*^(2)^.

**Figure 3:**
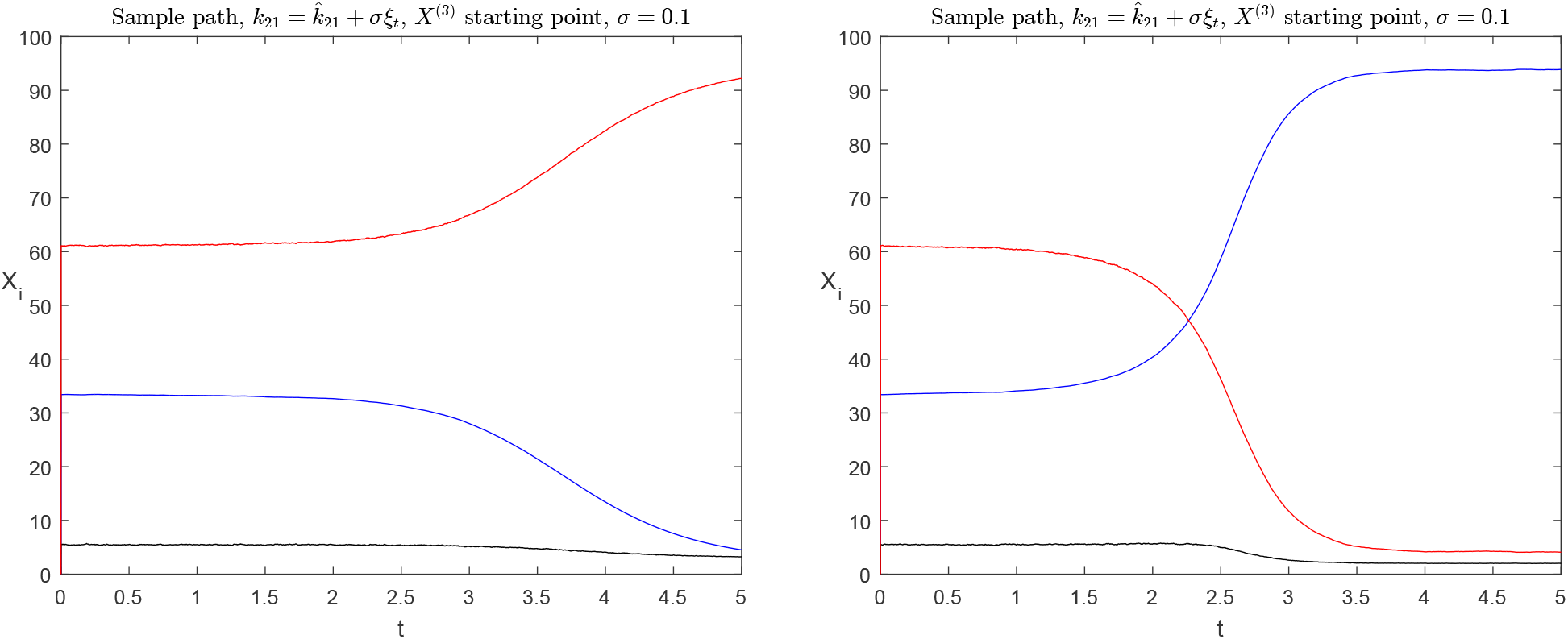
Sample paths of (3) with starting point *X*^(3)^ = (33.4, 5.5, 61.1)^*T*^ and *σ* = 0.1. Unmethylated CpG dyads are denoted in blue, hemimethylated in black and methylated in red. The sample paths converge either to the point *X*^(1)^ or *X*^(2)^.

**Figure 4:**
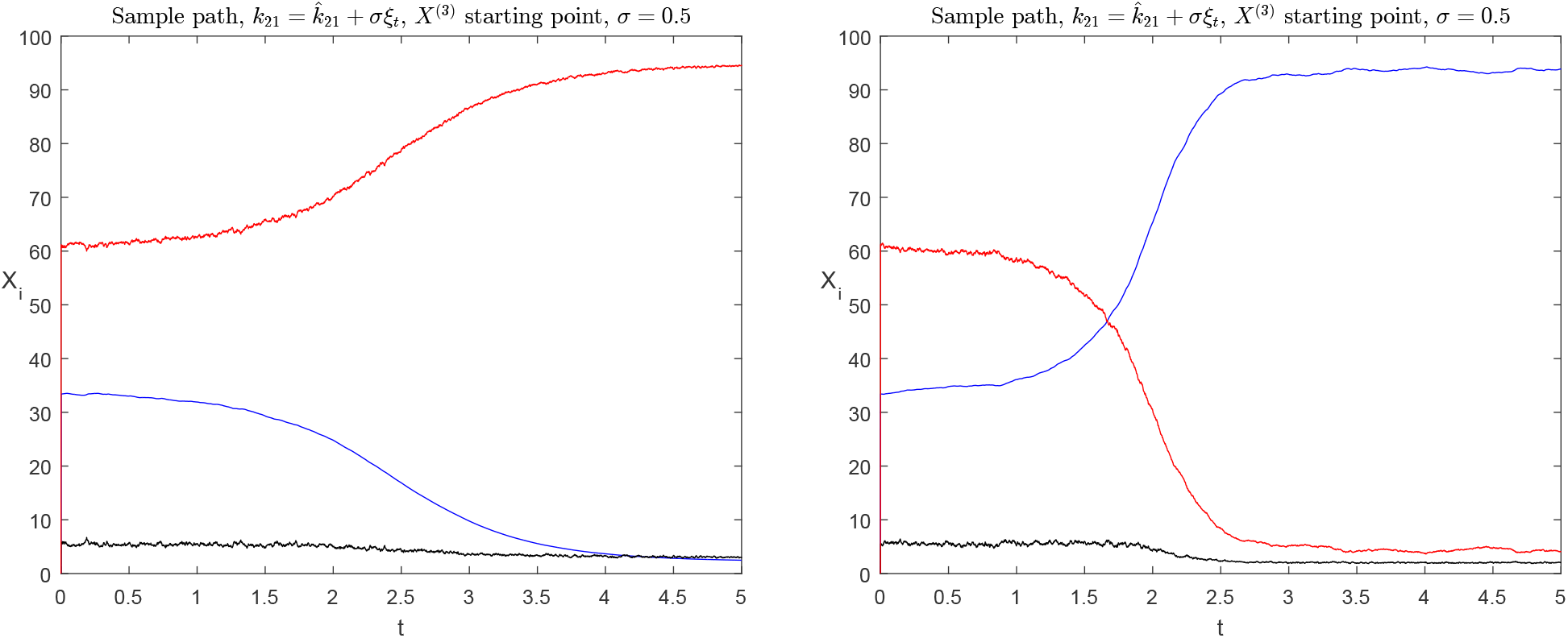
Sample paths of (3) with starting point *X*^(3)^ = (33.4, 5.5, 61.1)^*T*^ and *σ* = 0.5. Unmethylated CpG dyads are denoted in blue, hemimethylated in black and methylated in red. The sample paths converge either to *X*^(1)^ or *X*^(2)^.

**Figure 5:**
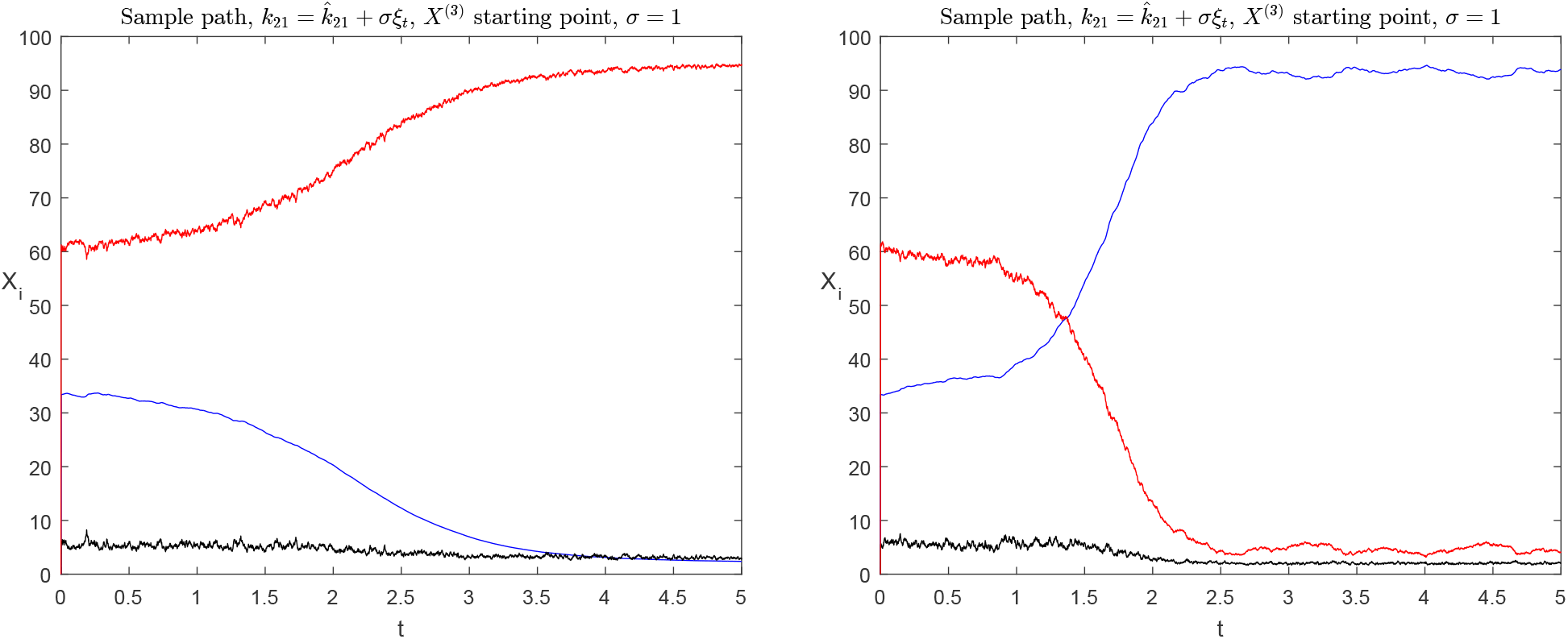
Sample paths of (3) with starting point *X*^(3)^ = (33.4, 5.5, 61.1)^*T*^ and *σ* = 1. Unmethylated CpG dyads are denoted in blue, hemimethylated in black and methylated in red. The sample paths converge either to *X*^(1)^ or *X*^(2)^.

Next we plot the mean trajectory of 10^4^ sample paths with noise level *σ* = 0.1, 0.5 and 1. Figure 6 shows that the value which the mean of 10^4^ sample paths converges to is different for different values of noise. The greater the noise level, the more unpredictable the convergence of the sample paths is.

**Figure 6:**
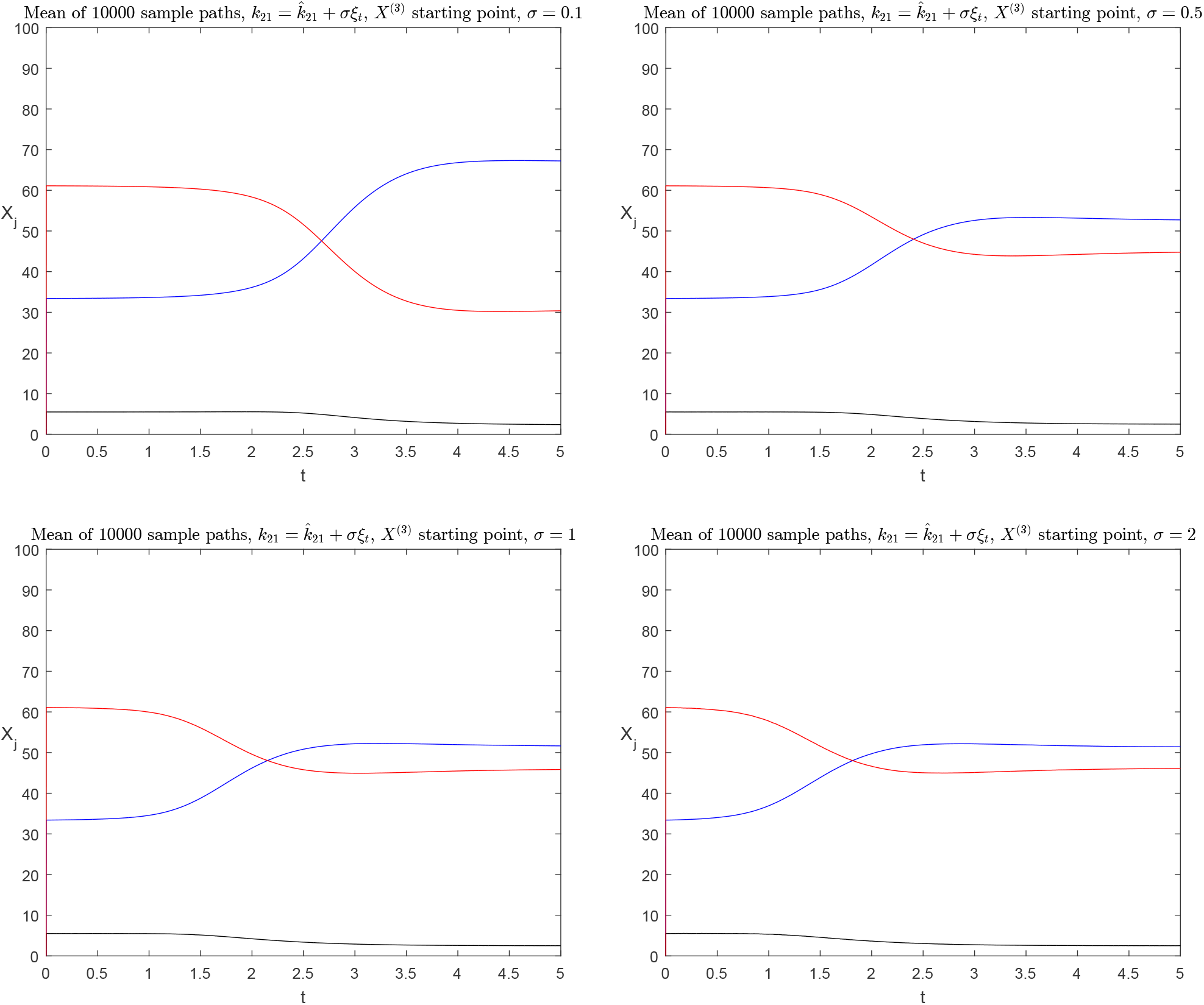
The mean of 10^4^ sample paths versus time. Unmethylated CpG dyads are denoted in blue, hemimethylated in black and methylated in red. All paths start from *X*^(3)^ = (33.4, 5.5, 61.1)^*T*^ and *σ* = 0.1, 0.5, 1, 2.

In other words, the closer to the boundary the initial value is, the higher the uncertainty to which steady point the stochastic trajectory converges. On the contrary, the closer to the steady points the initial value is, the more difficult it is for the sample paths to diverge from that steady point. After identifying that the points *X*^(1)^ = (2.2, 3.1, 94.7)^*T*^ and *X*^(2)^ = (93.2, 2, 4.1)^*T*^ are stable whereas point *X*^(3)^ = (33.4, 5.5, 61.1)^*T*^ is unstable, we need to define the area of uncertainty, where, if sample paths are plotted starting from the same initial point, some of them will converge to a different steady point than the rest. To do so, instead of plotting the trajectories of the solutions or mean trajectories, it is necessary to construct a grid of starting points.

To do so, a specific starting point *X*_0_ was selected and the ending point of a stochastic trajectory after *N* steps was identified. We calculated *M* stochastic trajectories starting from this initial value *X*_0_ and we identified the ending point of each one. If all *M* stochastic trajectories converge to steady point *X*^(1)^ = (2.2, 3.1, 94.7)^*T*^ then the starting point *X*_0_ was depicted in green. If all trajectories converge to steady point *X*^(2)^ = (93.2, 2, 4.1)^*T*^ then *X*_0_ was depicted in blue. Finally, if there is at least one stochastic trajectory converging to *X*^(1)^ and one converging to *X*^(2)^ then the initial value point was painted in red.

This process was repeated for various values of starting points *X*_0_ so that a grid of starting points was created, where green and blue colour denote convergence of all sample paths to *X*^(1)^ and *X*^(2)^, respectively, and red colour denotes uncertainty, namely, not knowing which steady point the sample paths will converge to. We constructed different grids for various values of noise *σ* and we investigated how increasing noise widens the region of uncertainty. The Euler-Maruyama scheme was used for approximating the solution of the stochastic differential equation (3).

This methodology was first implemented for the deterministic case, that is, *σ* = 0. As expected, there is a well defined boundary between the two regions of stability (Figure 7). All trajectories that start from the same initial point converge to the same point.

**Figure 7:**
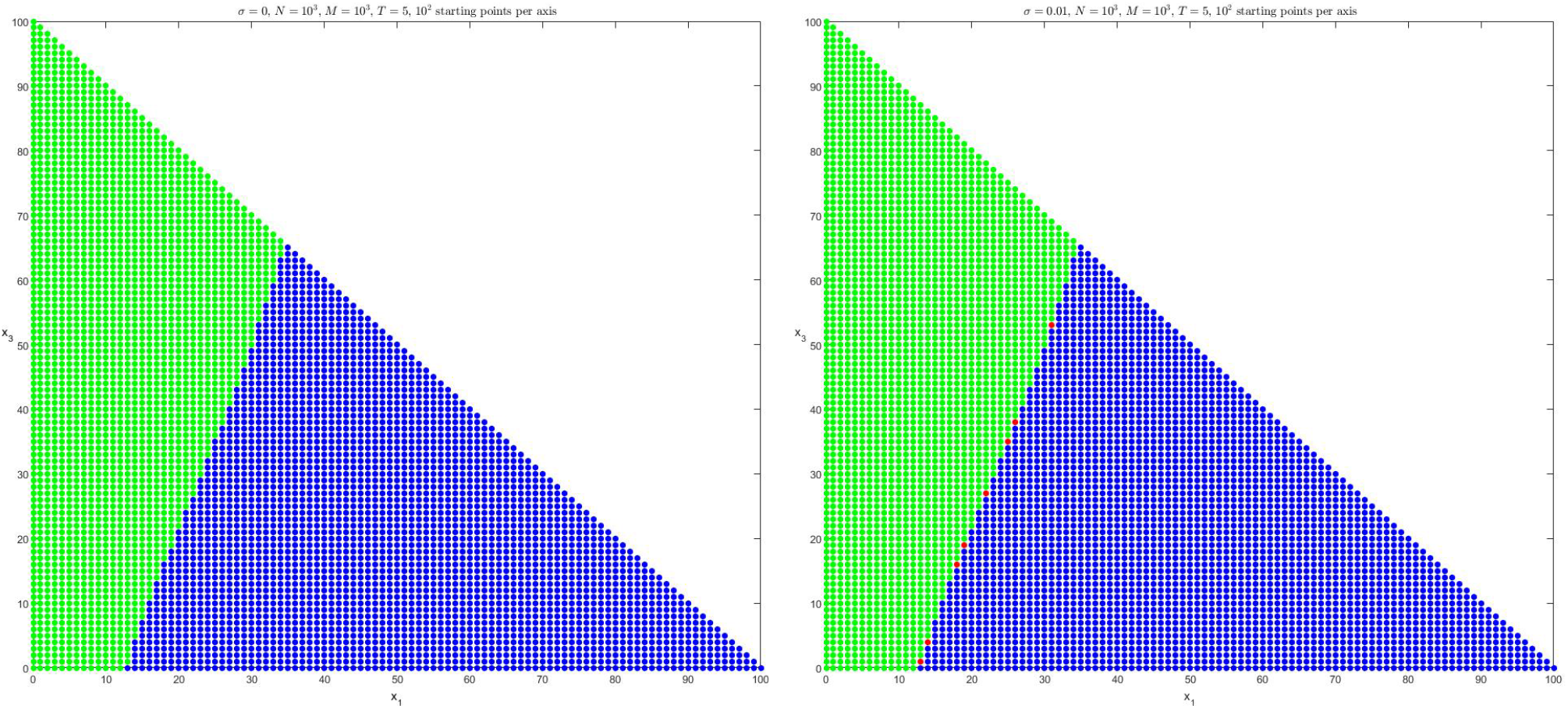
Grid of points in the *X*_1_ − *X*_3_ plane. On the left we have the deterministic case for *σ* = 0. On the right, *σ* = 0.01. Number of iteration steps *N* = 10^3^, final time *T* = 5, number of stochastic paths *M* = 10^3^ per initial point. Convergence of all stochastic trajectories to steady point *X*^(1)^ = (2.2, 3.1, 94.7)^*T*^ is denoted in green, to *X*^(2)^ = (93.2, 2, 4.1)^*T*^ in blue and convergence to both in red.

We added noise *σ* = 0.01 into the system. In the same figure, it can be seen that in the borderline of the two regions there are only a few sample paths that converge to different steady points, even though they start from the same point. Plotting the grids for the cases *σ* = 0.05, 0.1, 0.5, we see that the region of uncertainty grows as the noise level grows (Figures 8), which was well expected. Eventually, the region becomes quite large for *σ* = 1.

**Figure 8:**
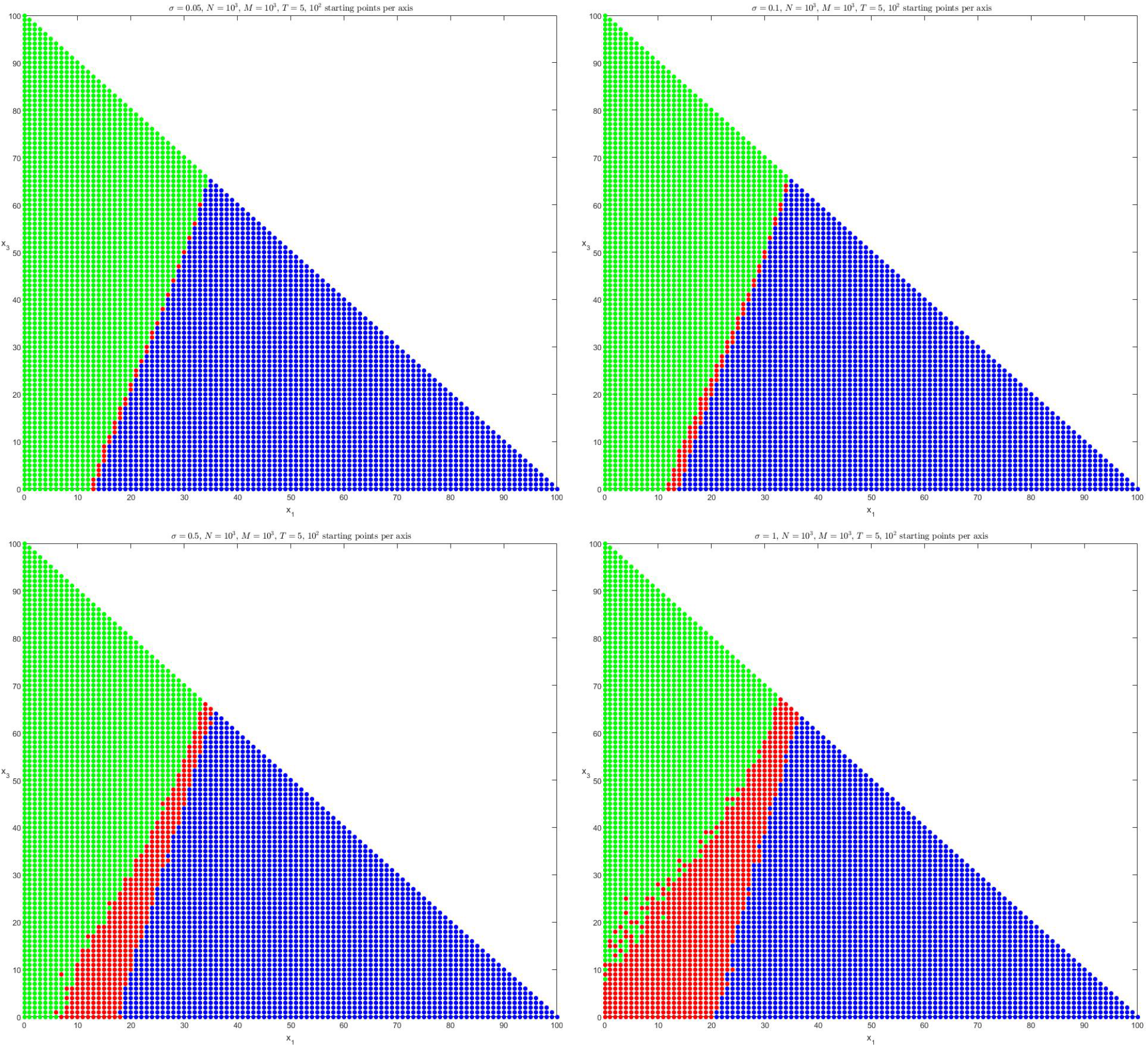
Grids of points in the *X*_1_ *− X*_3_ plane. Level of noise *σ* = 0.05, 0.1, 0.5, 1. Number of iteration steps *N* = 10^3^, final time *T* = 5, number of stochastic paths *M* = 10^3^ per initial point. Convergence of all stochastic trajectories to steady point *X*^(1)^ = (2.2, 3.1, 94.7)^*T*^ is denoted in green, to *X*^(2)^ = (93.2, 2, 4.1)^*T*^ in blue and convergence to both in red.

It is particularly interesting to note that uncertainty spreads towards both regions, but predominantly through the methylated region, especially for larger values of input noise. At high noise levels (50% − 100%), simulations show more cases where a convergence to high methylation levels (green) is replaced by the possibility of noisy divergence to low methylation levels (indicated by red). Since noise in parameter *k*_21_ primarily represents the stochastic change in maintenance methylation due to increasing DNA methylation age, it can be inferred that as methylation age progresses, CpG islands with intermediate methylation levels (the region close to the boundary in the simulations) tend to be more susceptible to dramatic DNA methylation changes towards both hypomethylation and hypermethylation. Methylation convergence uncertainty clearly spreads through the hypomethylated region (blue) as well, but seems to be less affected for the same amount of input noise.

## 4 Discussion

The ageing process is underpinned by a myriad of interacting biological mechanisms. Despite this complexity recent evidence has suggested, there could be certain general intracellular processes which characterise ageing. Among these mechanisms, changes to the DNA methylome have been acutely associated with the trajectory of the ageing process and how ageing intersects with healthspan. A particular area of intrigue surrounding the DNA methylome centres on the factors which precipitate a change in the methylation levels within gene promoters during ageing. Gene promoters have been found to be either in a hypomethylated or in a hypermethylated state, but rarely in an intermediate state [Haerter et al., 2014]. A change to the methylation level has the potential to impact genetic regulation which in turn could result in disease depending on the nature of the gene. In essence, if the gene promoter is hypomethylated, then transcriptional proteins are free to bind to the gene and thus the gene is active [Zilberman et al., 2007]. If the gene promoter is hypermethylated, then the methyl groups attached to the CpG sites impede the binding of transcriptional proteins to the gene, so that the gene is silenced [Razin, 1998]. If the gene is responsible for suppressing genetic mutations in the cell, its silencing would potentially lead to cancer development. Depending on the gene function, silencing due to hypermethylation could potentially lead to various age-onset pathologies [Baylin, 2005].

In this work we constructed a new stochastic mathematical model to investigate how age associated fluctuations in maintenance methylation influence promoter methylation. Randomness was introduced into the parameters of the model to account for the stochasticity in maintenance methylation. ’White noise’ was added and we examined how different levels of maintenance methylation noise impacted the model. We dynamically simulated stochastic trajectories using the stochastic model (3). Uncertainty was introduced to the most sensitive parameter of the model (1) and a new stochastic model (3) was created to account for the stochastic nature in cell dynamics. “White noise” was added to the process of maintenance methylation and it was investigated how various levels of input noise influence model outcomes.

In addition to *k*_21_ being the most sensitive parameter, there was also a clear biological rationale for examining this parameter. Noise in parameter *k*_21_ primarily represents the stochastic change in cite-specific DNA methylation due to increasing DNA methylation age. DNA methylation has been found to be an epigenetic clock which serves as a new standard to track chronological age and predict biological age [Xiao et al, 2019]. There is a clear association between increased DNA methylation age and risk of age-onset pathology [Fransquet et al., 2019]. Using the model we were able to quantify the uncertainty of cite-specific DNA methylation for different values of increasing age, which is represented by different values of input noise. Our model predicts the observed bistable levels in CpG islands, which is of key importance. In addition, simulations with various levels of noise indicate that uncertainty predominantly spreads through the methylated region, especially for larger values of input noise. This denotes that as age progresses, CpG islands with intermediate methylation levels tend to be more susceptible to dramatic DNA methylation level changes. Moreover, the closer the initial methylation levels are to the boundary between the hypomethylated and the hypermethylated regions, the more likely it is that the eventual methylation levels will alter significantly.

The limitations of the model are as follows. The validity of model findings depends on the validity of the biological assumptions made. Therefore, it is important to briefly review them. Even though gene promoter size may vary from gene to gene, an average sized gene promoter of a hundred CpG dyads length was adopted for the construction of the model. In addition, collaboration between neighbouring CpG sites was taken into consideration and therefore it was assumed that the transition rates vary over time as functions of the CpG population. This was a key assumption of the models which succeeds in portraying the bistable behaviour in gene promoters. Moreover, DNA methylation reactions were considered to be akin to second order kinetics and hence transition rate expressions were considered to be second order functions of the populations.

Another limitation centres on the estimation of parameter values, which was based solely upon previously published scientific papers. In order to estimate parameters using methylation data however, factors such as gender, age, ancestry and environmental factors such as diet, smoking and physical activity must be taken into consideration. This will result in a significantly more difficult model to construct with a more complex set of input parameters to be estimated, but will lead to a more valid representation of the process of DNA methylation. A further limitation of this work is that we only applied the model to a generic promoter with typical methylation levels. However, it is possible in the future to use the model to investigate genes that have been associated with ageing/healthspan. For instance, an updated stochastic model could potentially be applied to the study of a gene such as Adenomatous Polyposis Coli (APC). APC is the genetic blueprint which provides the instructions for synthesizing the protein APC; a large, multifunctional protein that is frequently mutated or down-regulated in colorectal, breast, renal, lung and prostate cancers [Kulis and Esteller, 2010] [Lesko et al., 2015]. Methylation data derived from studying the promoter of APC in tissues such as breast, kidney, colon and lung from healthy individuals of unknown age, gender and ancestry, suggest that 7% to 8% of the CpG dyads are methylated (Methbank database). However, 90 stage I lung cancer patients were tested for aberrant DNA methylation in five gene promoters [Harden et al., 2003] and APC showed the most frequent promoter hypermethylation (72% - 65 of 90). As an estimation of the levels of aberrant methylation in the APC promoter, data in lung tissue taken from a patient with adenocarcinoma UICC Stage III showed that methylation levels are almost at 100% (Methycancer database).

Therefore, if the exact sources of intracellular noise are identified it is possible to translate them into specific levels of ‘white noise’ in the model. Provided with this information we could then produce a *x*_1_ vs *x*_3_ plane using the stochastic methodology, which will pinpoint the methylation levels at which point there is a transition between a hypomethylated and hypermethylated states the in a the APC promoter. Such a finding could have significant implications for our overall understanding of how gene promoter methylation contributes to ageing, age related disease and healthspan.

## Acknowledgements

LZ would like to thank the University of Chester for funding his PhD studentship.

## Appendix

## Wiener Process

Based on [Kloeden and Platen, 1999], we define a standard Wiener process *W* = {*W*_*t*_, *t* ≥ 0} to be a Gaussian process with independent increments such that

- *W* (0) = 0, with probability one.
- *E*(*W*(*t*)) = 0
- Var(*W*(*t*) − *W*(*s*)) = *t* − *s*

for all 0 ≤ *s* ≤ *t*.

## White Noise

Assume the stochastic differential equation

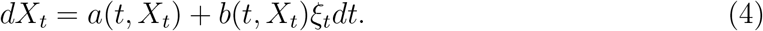

Here, a deterministic drift term *a*(*t, X*_*t*_) is perturbed by a noisy, diffusion term *b*(*t, X*_*t*_)ξ_*t*_, where ξ_*t*_ is a white noise process (standard Gaussian random variable) for each *t* and *b*(*t, X_t_*) a space-time dependent intensity factor. The stochastic differential equation 4 can be interpreted as an integral equation

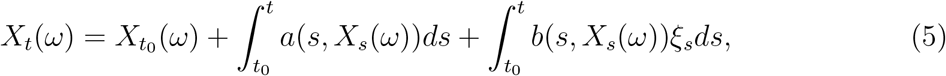

for each sample path. ξ_*t*_ can be seen as the generalised derivative of pure Brownian motion, that is, the derivative of a Wiener process *W*_*t*_, thus suggesting that we could write equation (5) as follows

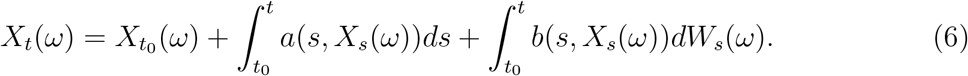

## Derivation of the stochastic model

We introduce randomness into parameter *k*_21_ as follows

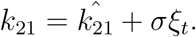

We rewrite the nonlinear system (1) with respect to *k*_21_

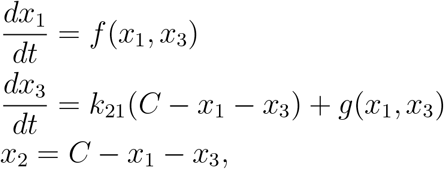

where

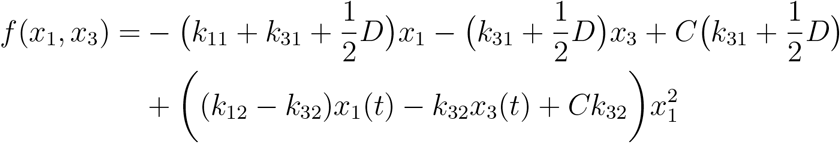

and 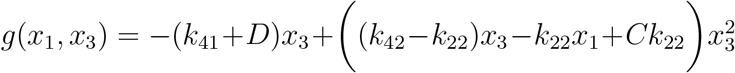. We substitute the expression (2) for the parameter *k*_21_ into the above system and obtain:

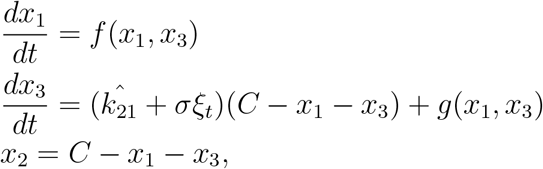

or rearranged

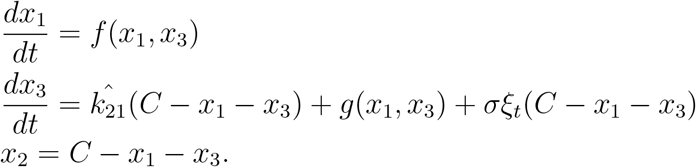

Since noise was added to one parameter of the system, the above system of ordinary differential equations turns into a system of stochastic ordinary differential equations. It would be better at this point to represent the equations in a more appropriate way, as follows:

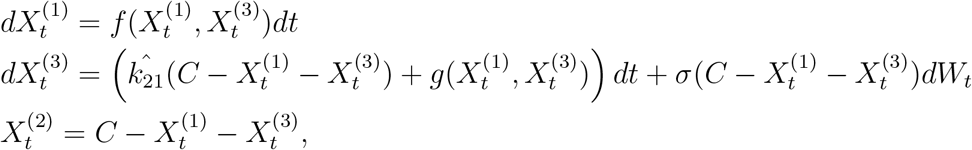

where 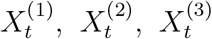 are now random variables and *W*_*t*_ is a scalar standard Wiener process with independent components associated with an increasing family of *σ*-algebras 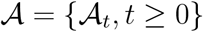. Equivalently, the above system can be denoted in matrix form, as

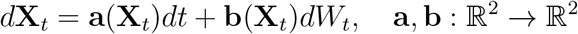

where

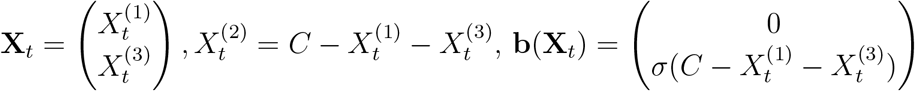

and

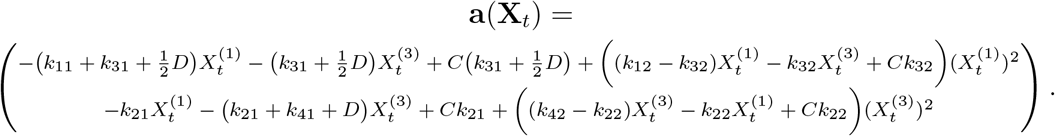

For the value of the parameter 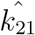, we used the nominal value of the parameter *k*_21_ used in the deterministic model. Thus, for simplicity reasons, we substituted the term 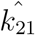 with *k*_21_. Equivalently, the system can be denoted with stochastic integral equations as follows

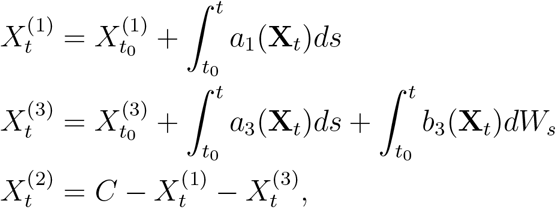

where

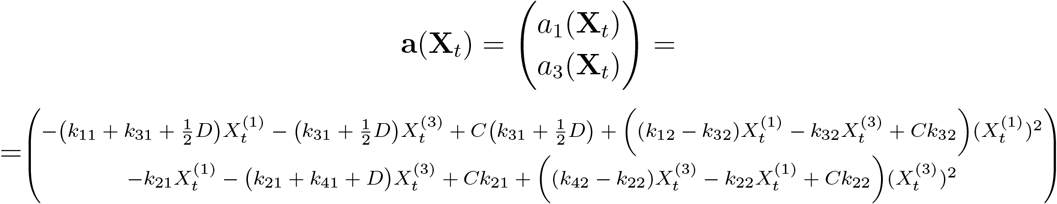

and

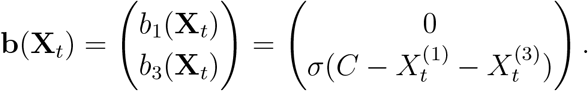

## Supplementary Materials

### Figures

**Figure 9:**
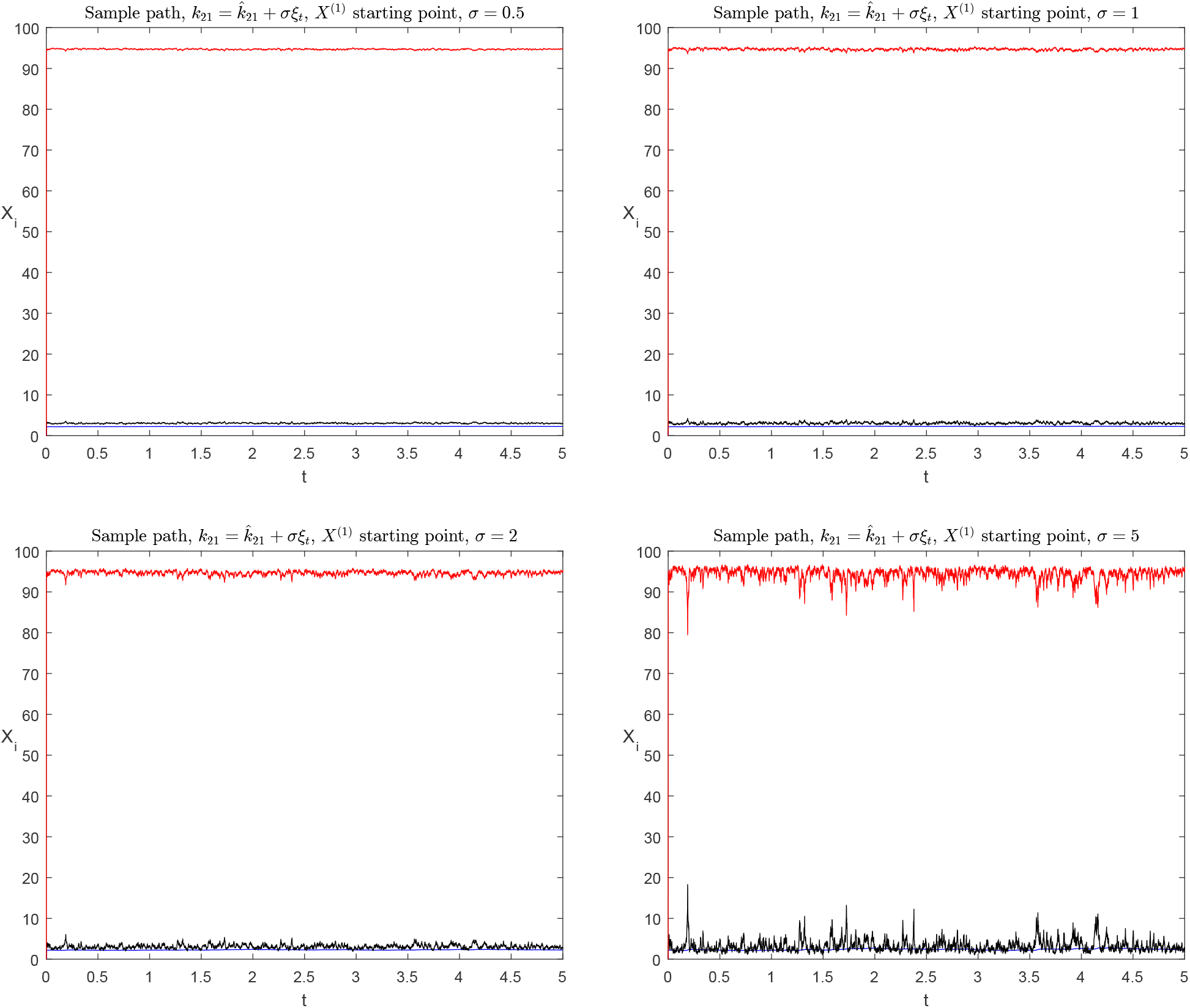
A sample path of (3) with starting point *X*^(1)^ = (2.2, 3.1, 94.7)^*T*^. Unmethylated CpG dyads are denoted in blue, hemimethylated in black and methylated in red. Below *σ* = 0.5, sample paths appear similar to deterministic solutions, having negligible fluctuation.

**Figure 10:**
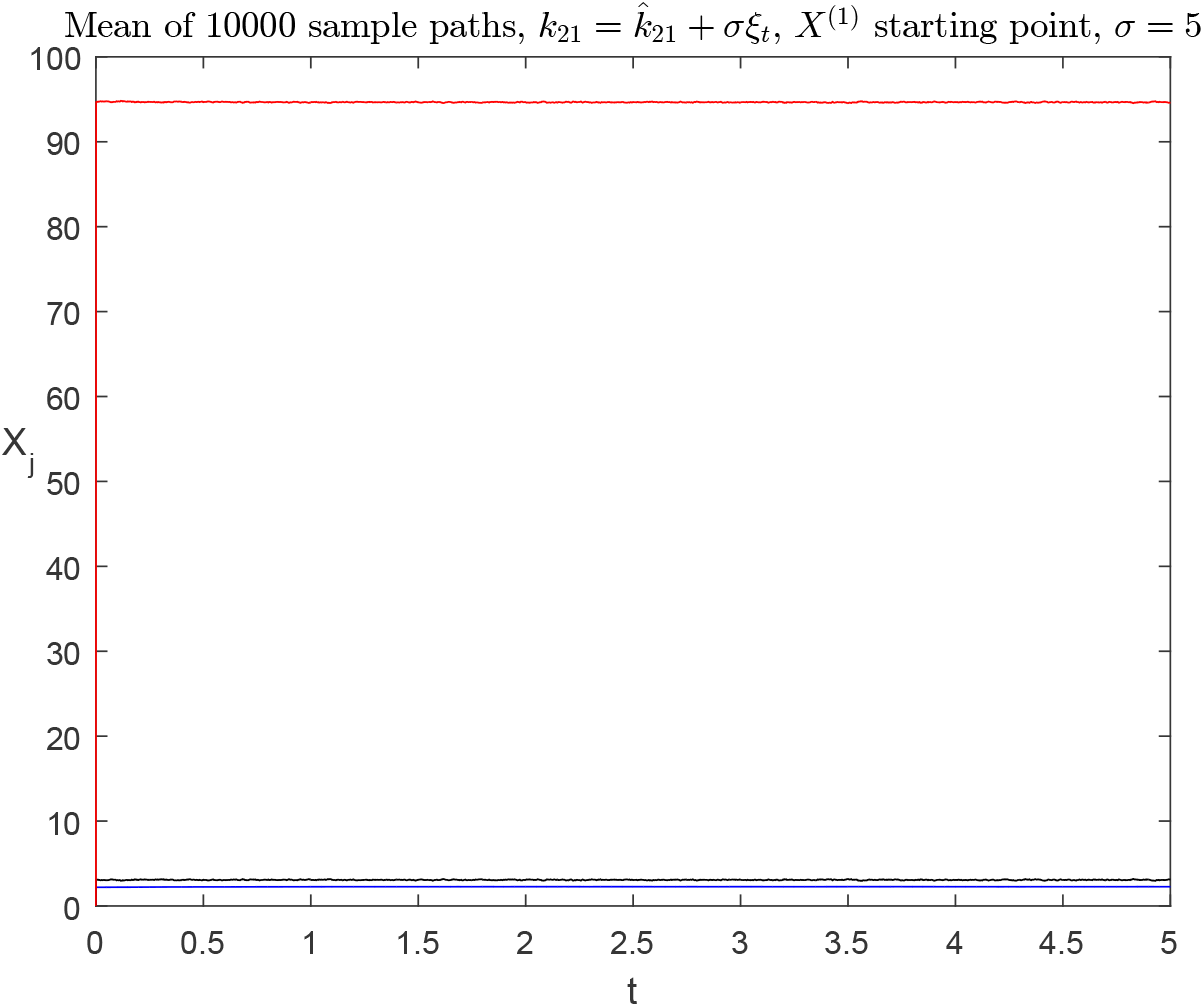
The mean of 10^4^ sample paths versus time. Unmethylated CpG dyads are denoted in blue, hemimethylated in black and methylated in red. All paths start from *X*^(1)^ = (2.2, 3.1, 94.7)^*T*^. The mean converges to *X*^(1)^.

**Figure 11:**
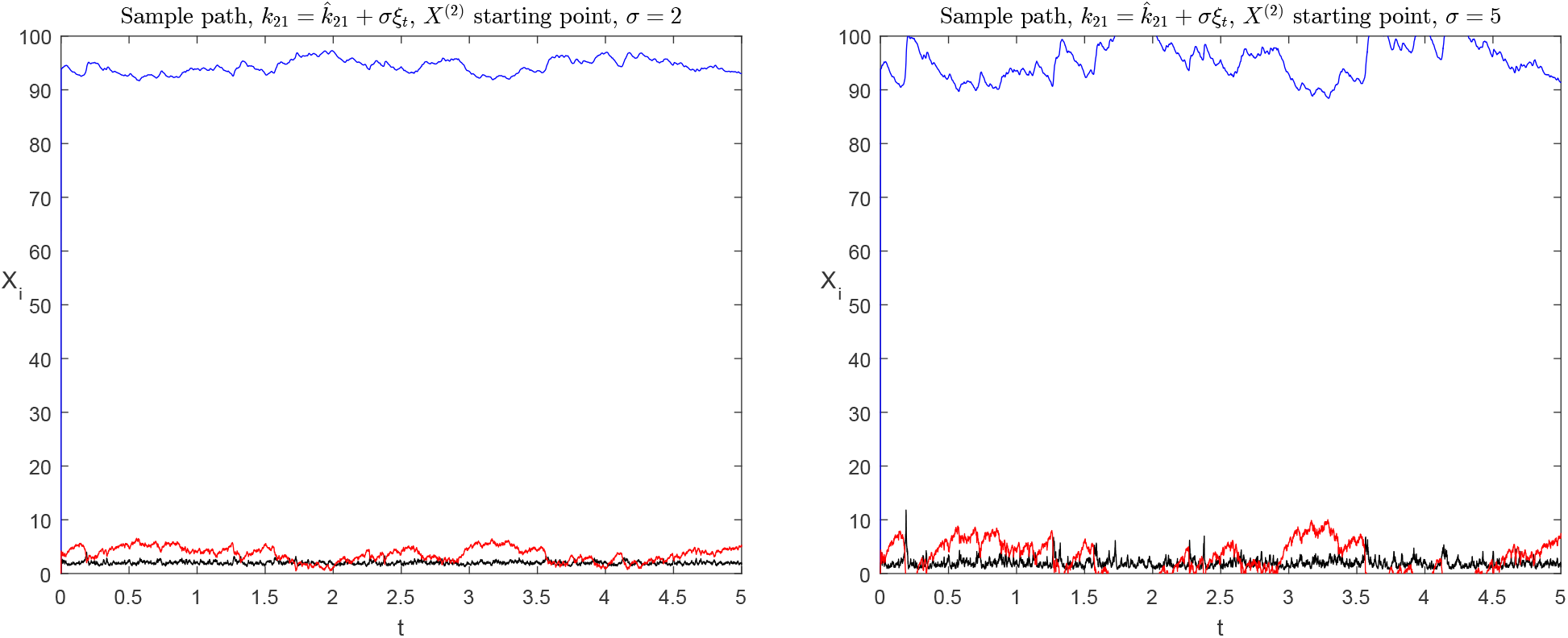
A sample path of (3) with starting point *X*^(2)^ = (93.2, 2, 4.1)^*T*^ with input noise *σ* = 200% and *σ* = 500%. Unmethylated CpG dyads are denoted in blue, hemimethylated in black and methylated in red. The sample paths have more significant fluctuations near the point *X*^(2)^ than near *X*^(1)^.

**Figure 12:**
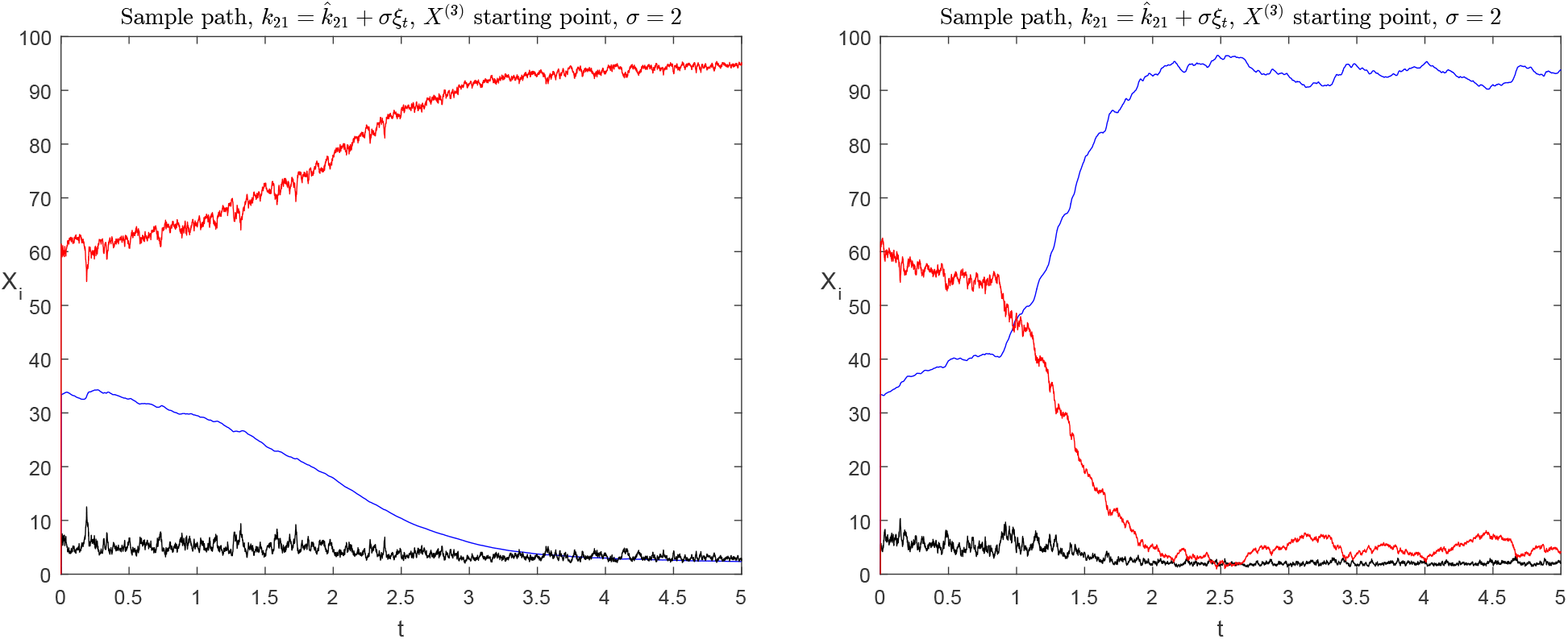
Sample paths of (3) with starting point *X*^(3)^ = (33.4, 5.5, 61.1)^*T*^ and *σ* = 2. Unmethylated CpG dyads are denoted in blue, hemimethylated in black and methylated in red. The sample paths converge either to *X*^(1)^ or *X*^(2)^.

